# The Ran GTPase Gradient Protects the Nucleolus from Aging-Associated Morphological Changes

**DOI:** 10.1101/2020.07.24.220079

**Authors:** Bartlomiej Remlein, Bryce M. Paschal

**Affiliations:** Center for Cell Signaling, University of Virginia; Department of Biochemistry and Molecular Genetics, University of Virginia

**Keywords:** Ran GTPase, nuclear transport, nucleolus, senescence, aging, nucleolin, B23/nucleophosmin, chromatin

## Abstract

In the context of its regulatory function for nucleocytoplasmic transport, the Ran GTPase undergoes cycles of nuclear import, GTP loading, nuclear export, and GTP hydrolysis. These reactions give rise to a nuclear:cytoplasmic (N:C) Ran gradient. In Hutchinson-Gilford Progeria Syndrome, disruption of the Ran gradient suppresses nuclear import of high molecular mass complexes by reducing the nuclear concentration of Ran. Here, we report that cells undergoing senescence, as a consequence of passage number, chemical induction, and altered nuclear lamina structure, all display a Ran gradient disruption quantitatively similar to that observed in Progeria patient cells. We found that the Ran gradient is critical for maintenance of nucleolar structure, as its disruption increases the size and decreases the average number of nucleoli per cell. Nucleolar number and size are biomarkers of longevity in diverse organisms, thus the nuclear level of Ran may be important for the nucleolar morphology in aging. The contribution of the Ran gradient includes regulating import of nucleolin and nucleophosmin, nucleolar proteins that assemble into high molecular mass complexes. The steepness of the Ran gradient is highly dependent on nuclear heterochromatin, which is reduced by passage number and chemical induction of senescence in cultured cells, and is known to decline during normal aging. Our data suggest that the Ran gradient senses nuclear heterochromatin, and through its function as a transport regulator, helps maintain the protein composition and structure of the nucleolus.

## INTRODUCTION

The nuclear transport machinery mediates the movement of proteins and RNA through nuclear pore complexes (NPCs) embedded in the nuclear envelope. Import and export pathways involve signal recognition by transport receptors, and translocation through the NPC facilitated by transient interactions with nucleoporins that line the NPC channel (Kim et al., 2018). Regulation of import and export pathways in all eukaryotes is provided by the Ran GTPase (Gorlich & Kutay, 1999). Nuclear transport regulation by Ran is dependent on a GTPase cycle and a nucleocytoplasmic transport cycle, both of which are important for its role in controlling transport complex assembly and disassembly (Dasso & Pu, 1998; Pemberton & Paschal, 2005; Stewart, 2007). The key steps of GTP loading and GTP hydrolysis are compartmentalized by virtue of the mutually exclusive localization of RCC1, the RanGEF (nucleus), and RanGAP (cytoplasm). Ran is mostly nuclear under steady state conditions, but Ran export from the nucleus occurs as a continuous process as it exits the nucleus bound to transport receptors. Ran reentry into the nucleus is mediated by NTF2, a factor dedicated to nuclear import of Ran-GDP (Ribbeck, Lipowsky, Kent, Stewart, & Gorlich, 1998; Smith, Brownawell, & Macara, 1998; Steggerda, Black, & Paschal, 2000).

The integration of GTPase and nucleocytoplasmic transport cycles results in a nuclear:cytoplasmic (N:C) Ran protein gradient, which is approximately 3:1 when measured by fluorescence microscopy in mammalian cells and in budding yeast (Kelley & Paschal, 2019). The Ran gradient is disrupted in cells from Hutchinson-Gilford Progeria Syndrome (HGPS) patients, which we showed is caused specifically by constitutive attachment of laminA to the nuclear membrane (Kelley et al., 2011). Preventing lamin A farnesylation, which mediates lamin A attachment to the inner nuclear membrane, rescues the Ran gradient (Kelley et al., 2011). For reasons that are still not understood, constitutive nuclear membrane anchoring of lamin A is sufficient to cause a reduction in nuclear heterochromatin in HGPS (Scaffidi & Misteli, 2006; Shumaker et al., 2006). Because RanGEF shows a preferential association with heterochromatin, and chromatin provides a scaffold for GTP loading by the RanGEF, we proposed that Ran gradient disruption in HGPS cells is a consequence of altered lamina structure and reduced heterochromatin that lowers Ran-GTP production (Dworak et al., 2019). Consistent with this model, pharmacological inhibition of the histone methyltransferase G9a is sufficient to disrupt the Ran gradient (Dworak et al., 2019).

Reduced levels of heterochromatin is one of the hallmarks of normal aging (Lopez-Otin, Blasco, Partridge, Serrano, & Kroemer, 2013), which is a part of the evidence that there are cellular features shared between premature and normal aging. In the current study, we set out to address whether the Ran gradient disruption observed in HGPS occurs in other aging-related settings. We determined that the Ran gradient is disrupted in cells treated with tert-butyl hydroperoxide to reach senescence, as well as in cells allowed to undergo replicative senescence. We found that Ran gradient disruption can account for aging-associated phenotypes in the nucleolus. This involves a Ran gradient-dependent import of nucleolin and nucleophosmin, proteins with multiple nuclear functions that includes regulation of nucleolar structure and output. Our data suggests the Ran gradient plays an important role in sensing and transducing cellular aging phenotypes.

## RESULTS

Our previous analysis of HGPS cells showed that the structure of the nuclear lamina is functionally linked to heterochromatin and the Ran gradient; these relationships define an axis that upon disruption results in a reduction in nuclear import (Dworak et al., 2019; Kelley et al., 2011; Snow, Dar, Dutta, Kehlenbach, & Paschal, 2013). In a variety of species, heterochromatin undergoes age-dependent reduction and redistribution (Tsurumi & Li, 2012; Villeponteau, 1997). We posited that heterochromatin reduction might provide a mechanism for controlling the Ran gradient and nuclear transport in senescence and aging. By this logic, a declining level of heterochromatin characteristic of aging cells is predicted to reduce nuclear transport of high molecular weight (> 300 kD) proteins and complexes, which in HGPS cells (Snow et al., 2013) are preferentially affected by changes in Ran distribution.

To determine if senescence affects the Ran pathway, we adopted a protocol for senescence induction (Debacq-Chainiaux, Erusalimsky, Campisi, & Toussaint, 2009; Dumont et al., 2000) that involved sequential exposure of low passage, normal human fibroblasts to tert-butyl hydroperoxide (tBHP), followed by a 2-day period for recovery from any acute effects of the treatment (**Fig. S1A**). Human fibroblasts exposed to tBHP showed an induction of senescence-associated β-galactosidase (SA-β-gal), and by immunofluorescence (IF) microscopy, a significant reduction in the N:C Ran gradient (**Fig. 1A-C**; p < 0.001). From double label-IF for Ran and Histone H3K9me3, we found that cells with low levels of heterochromatin had a reduced Ran gradient (p < 0.001; **Fig. 1D, E**). A comparison of early and late passage fibroblasts (P11 vs. P23) showed the anticipated increase in senescent cells, with late passage cells displaying the same changes in heterochromatin and the Ran gradient observed after tBHP treatment (**Fig. 1F-J**). To explore whether there is a causal relationship between the Ran gradient and senescence, we used siRNA to deplete the Ran import factor NTF2, and examined the cells by microscopy. Disruption of the Ran gradient by NTF2 depletion was sufficient to induce senescence and drive a reduction in heterochromatin (**Fig. 1K-O**). Comparable effects were also obtained by treating fibroblasts with lopinavir (LPV), which induces a progeric phenotype by inhibiting pre-lamin A processing (**Fig. 1P-T**). These data show that in senescent cells, there is a disruption of the Ran GTPase gradient. Since reducing Ran import into the nucleus is sufficient to trigger senescence, the Ran gradient appears to be important for preventing cells from entering a senescent state. Thus, the Ran gradient appears to have a geroprotective effect on the cell.

**Fig. 1.**
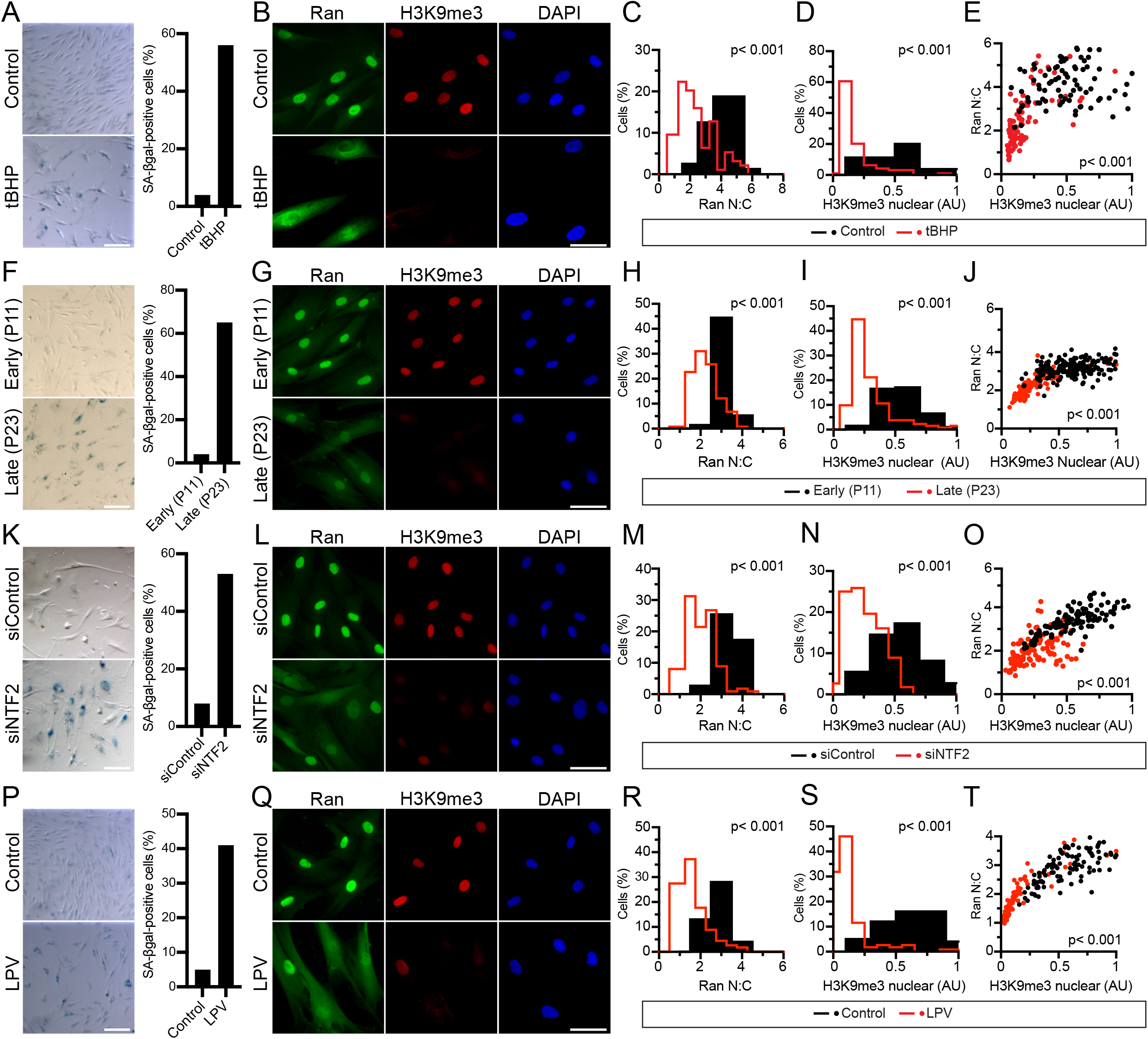
The Ran gradient is disrupted in senescence. **(A, F, K, P)** Phase contrast images of SA-β-gal assays performed on normal human fibroblasts exposed to tert-butyl hydroperoxide (tBHP), passaged for replicative senescence, depleted of NTF2, and treated with lopinavir (LPV), respectively. Scale bar = 150 μm. Percentage of β-gal positive cells scored and plotted (n = 150). **(B, G, L, Q)** Double label IF microscopy showing the distribution of Ran and histone H3K9me3. Scale bar = 60 μm. (**C, D, H, I, M, N, R, S)** Quantification of IF microscopy showing Ran N:C and H3K9me3 nuclear fluorescence (n = 100). **(E, J, O, T)** Plots of Ran N:C versus H3K9me3 nuclear fluorescence from individual cells (n = 100). p values are indicated in the panels.

Aging and longevity pathways converge on the nucleolus via mechanisms that help coordinate ribosome biogenesis with nutrient sensing (Tiku & Antebi, 2018). Pioneering studies in model organisms and subsequent work in mammalian cells led to the view that small nucleoli are associated with longevity, and that enlargement of nucleoli occurs during both normal and premature aging (Buchwalter & Hetzer, 2017; Tiku et al., 2017). Using acetyltransferase NAT10 as an endogenous nucleolar marker (Ito et al., 2014), we found that senescence induction with tBHP caused nucleolar enlargement, resulting in a significant increase in the proportion of cells with 1-2 nuclei, though interestingly, the total nucleolar area per nucleus was affected only slightly (**Fig. 2A, B, C, D**). These nucleolar changes were also observed when the Ran gradient was disrupted by NTF2 depletion, as well as by treatment with LPV (**Fig. 2A, B, C; Fig. S1B-I**). Nucleoli in senescent cells and in NTF2 depleted cells retained fibrillarin positivity (**Fig. 2A, H; Fig. S1B, F**). There was, however, a marked redistribution of nucleolin from nucleoli to the cytoplasm of cells treated with tBHP, siNTF2, and LPV, which occurred without changes in nucleolin protein levels detected by immunoblotting (**Fig. 2E, F**). Nucleolin shuttles between the nucleus and cytoplasm (Borer, Lehner, Eppenberger, & Nigg, 1989), but it is highly concentrated in the nucleolus under steady state conditions, and the small cytoplasmic pool is not readily seen by IF microscopy (**Fig. 2E**). These data indicate that nucleolin import into the nucleus is highly reduced in senescent cells under conditions where there are negligible changes in nuclear import and nucleolar localization of NAT10 and fibrillarin (**Fig. 2G, H, I**). NAT10 was reported by another group to have nuclear import defects in HGPS cells (Larrieu et al., 2018), but in our assays NAT10 is efficiently localized to nucleoli even when the Ran gradient is disrupted.

**Fig. 2.**
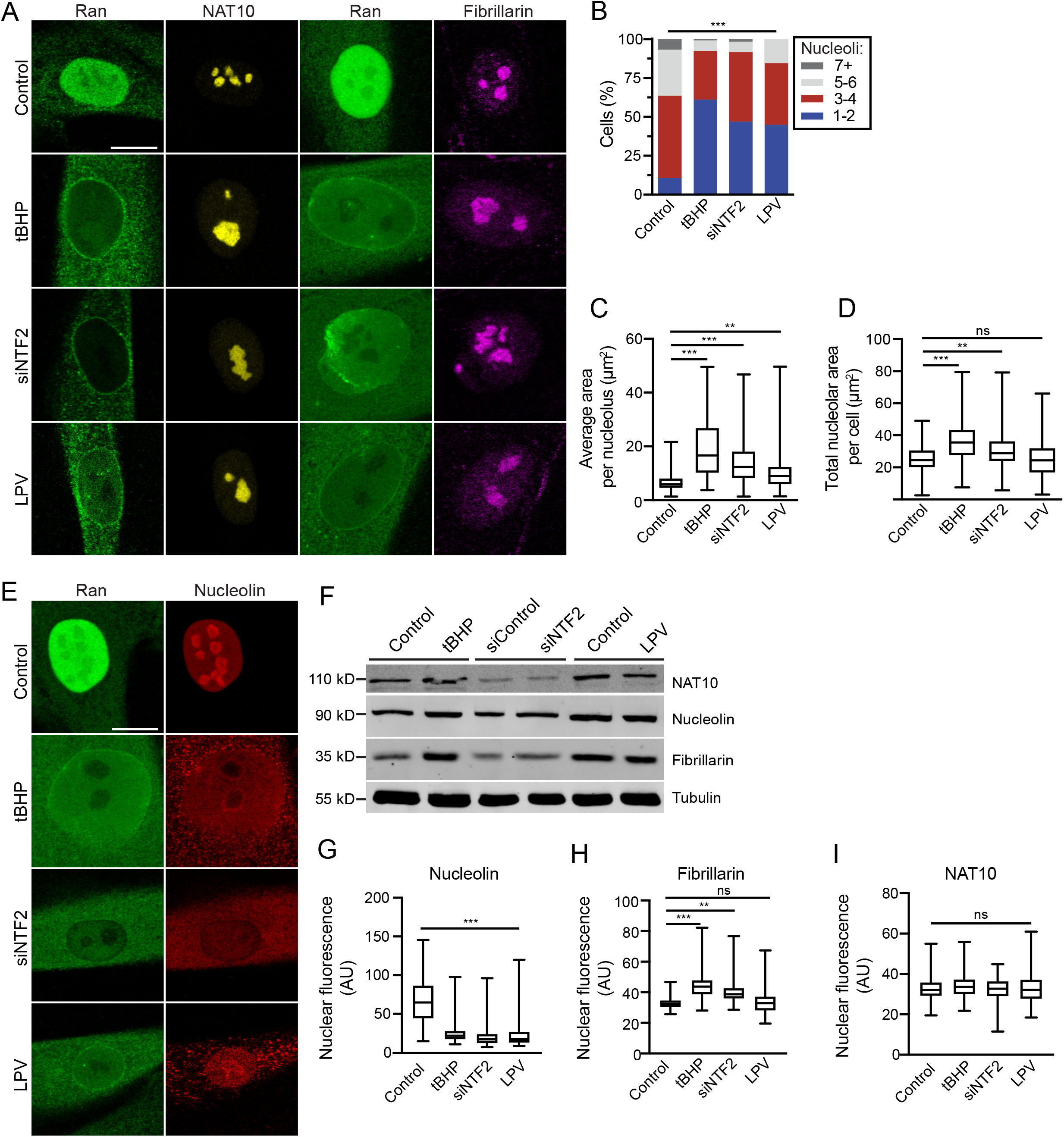
Nucleolar structure is dependent on the Ran gradient. **(A)** Double label IF microscopy showing distribution of ran, NAT10 and fibrillarin in normal human fibroblasts exposed to tBHP, depleted of NTF2, and treated with LPV. Scale bar = 10 μm. **(B)** Number of nucleoli per cell quantified using NAT10 staining (n = 121). **(C)** Box and whiskers plot of average nucleolar size (n = 121). **(D)** Box and whiskers plot of total nucleolar size (n = 121). **(E)** Double label IF microscopy showing distribution of ran and nucleolin in normal human fibroblasts exposed to tBHP, depleted of NTF2, and treated with LPV. Scale bar = 10 μm. **(F)** Western blotting for NAT10, nucleolin, fibrillarin, and tubulin (loading control) under indicated conditions. **(G, H, I)** Box and whiskers plot of total nuclear fluorescence of nucleolin, fibrillarin and NAT10 under indicated conditions. (n = 150). In plots, boxes denote 25^th^ and 75^th^ percentile with median in the middle, and whiskers show minimal and maximal values. (ns, not significant, p > 0.05; *, p < 0.05; **, p < 0.001; ***, p < 0.0001, ANOVA).

Nucleophosmin/B23 (NPM), like nucleolin, is an abundant, multi-functional, nucleolar protein that participates in ribosome biogenesis (Colombo, Alcalay, & Pelicci, 2011), and over-expression of NPM can suppress oncogene-induced senescence (Li, Sejas, Burma, Chen, & Pang, 2007). We examined NPM localization and found both its nucleolar and nuclear localization were dramatically lower in cells treated with tBHP and LPV, as well as in cells depleted of the Ran import factor NTF2 (**Fig. 3A**). Thus, nuclear import of NPM is reduced by conditions that promote senescence and disrupt the Ran gradient. Because nuclear import of high molecular weight proteins and complexes are sensitive to perturbation of the Ran gradient (Dworak et al., 2019; Snow et al., 2013), we examined the sizes of endogenous nucleolin and NPM in cell extract using gel filtration chromatography. NPM eluted with an apparent mass of ~614 kD, while the size of nucleolin determined by this method was ~336 kD (**Fig. 3B, C**). Given these masses are much larger than the sizes of nucleolin (710 aa) and NPM (294 aa), it is clear these proteins exist as large protein complexes. We focused on nucleolin transport to formally test the hypothesis that its localization defect in response to senescence and Ran disruption is due to its size, as opposed to inactivation of the NLS. For this analysis, we fused the nucleolin NLS to inert proteins (**Fig. 3D, E, G)** and generated intermediate (60 kD; double GFP) and large (336 kD; pyruvate kinase tetramer) reporter proteins (Snow et al., 2013). To disrupt the Ran gradient, we co-transfected HA-Progerin with the reporters, and as a control, a HA-Progerin (C611S) farnesylation deficient mutant that does not undergo membrane anchoring, and therefore, does not affect Ran distribution (Snow et al., 2013). Nuclear import mediated by the nucleolin NLS fused to the large reporter protein (myc-PK-nucleolin NLS) was inhibited by Progerin, but its nuclear import was unaffected when co-expressed with the farnesylation-defective Progerin (**Fig. 3E, F**). By contrast, nuclear import mediated by the intermediate sized reporter protein (GFP-GFP-nucleolin NLS) was not inhibited by Progerin despite the fact that the Ran gradient was clearly disrupted (**Fig. 3G, H**). While it is certainly possible that additional senescence-related mechanisms could affect nuclear localization of nucleolin, our data shows that the molecular size of nucleolin is a critical feature that dictates its import efficiency in cells with Ran system defects.

**Fig. 3.**
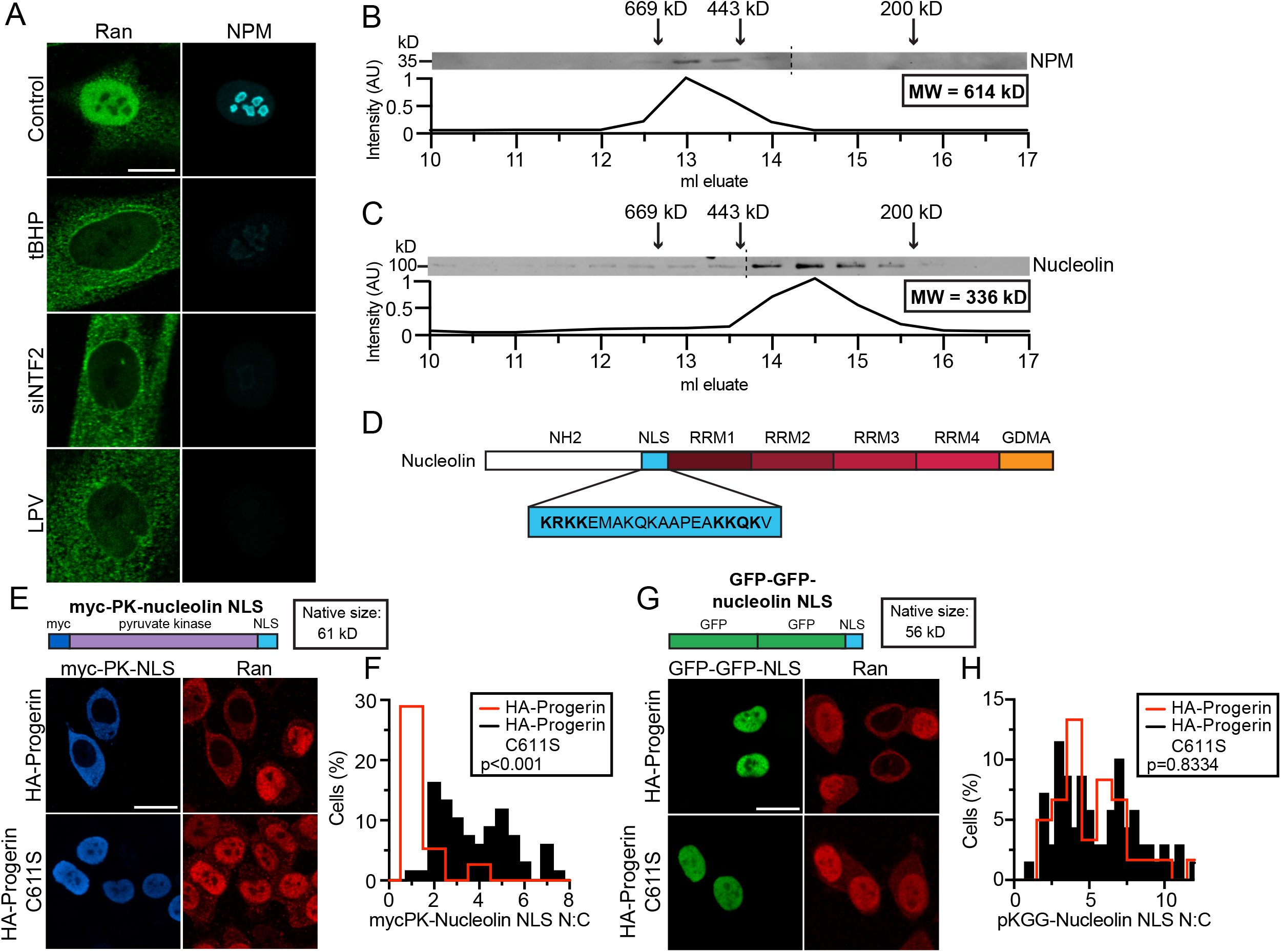
Nucleolar proteins nucleolin and nucleophosmin form high molecular weight complexes and undergo Ran gradient-dependent import. **(A)** Double label IF microscopy showing distribution of ran and nucleophosmin in normal human fibroblasts exposed to tBHP, depleted of NTF2, and treated with LPV. Scale bar = 10 μm. (**B-C**) Gel filtration chromatography of 293T cell lysates showing the elution positions of nucleophosmin and nucleolin by immunoblotting and plotted as band intensity. Positions of molecular standards are indicated by arrows. **(D)** Domain structure of nucleolin showing the position of bipartite nuclear localization signal (NLS) in blue. **(E-H)** Reporter protein fusions for analysis of nucleolin import. **(E)** Pyruvate kinase fusion (61 kD) fused to the nucleolin NLS and co-expressed with HA-progerin and farnesylation mutant of HA-progerin. Double label IF microscopy of myc-PK-Nucleolin-NLS and endogenous ran. Scale bar = 20 μm. **(F)** Quantification of IF microscopy showing myc-PK-nucleolin NLS N:C ratio signal (n = 70). **(G)** GFP-GFP fusion (56 kD) fused to the Nucleolin NLS and coexpressed with HA-Progerin and farnesylation mutant of HA-Progerin. Double label IF microscopy of GFP-GFP-Nucleolin-NLS and endogenous Ran. Scale bar = 20 μm. **(H)** Quantification of IF microscopy showing GFP-GFP-Nucleolin NLS N:C ratio signal (n = 70).

Both nucleolin and NPM function on multiple nuclear pathways, most notably the regulation of events that impact ribosome biogenesis (Pederson, 2011). We tested whether depletion of nucleolin and NPM is sufficient to generate the nucleolar changes observed in senescent cells. Using siRNA, we found that modest depletion of these two proteins individually was sufficient to affect nucleolar morphology (**Fig. 4A, E**). Nucleolin depletion reduced the number of nucleoli per cell and increased the average area per nucleolus with a penetrance similar to that observed with NTF2 depletion (**Fig. 4B, C**). Reducing NPM levels resulted in a subset of nucleoli with highly irregular shapes, which has been reported (Amin, Matsunaga, Uchiyama, & Fukui, 2008), but NPM depletion did not significantly change the number of nucleoli, or the average area per nucleolus (**Fig. 4C, D**). Conditions that promote enlargement of nucleoli can increase ribosome synthesis and protein translation (Buchwalter & Hetzer, 2017). To determine if senescence induction by tBHP, and manipulation of NTF2, nucleolin, and NPM levels affects translational output in cells, we used the SUnSET assay with an antibody that detects puromycin incorporation in proteins; puromycin can act as a mimetic of aminoacyl-tRNA (Schmidt, Clavarino, Ceppi, & Pierre, 2009). Immunofluorescence microscopy and quantification of puromycin staining showed that its incorporation into proteins was elevated in senescent fibroblasts generated by tBHP treatment, and as expected, labeling was blocked by cycloheximide (**Fig. 4F**). In tBHP-treated cells, there was a higher level of puromycin labeling indicating an increase in translation, though interestingly, the differences were still detected when cells were binned on the basis of nucleoli number (**Fig. 4G;** p < 0.001). Puromycin labeling was increased in cells depleted of NTF2, and to a lesser extent, by depletion of nucleolin and NPM (**Fig. 4H;** p < 0.001). These data show that the status of the Ran gradient and nucleolar protein composition has a quantitative effect on translational output.

**Fig. 4.**
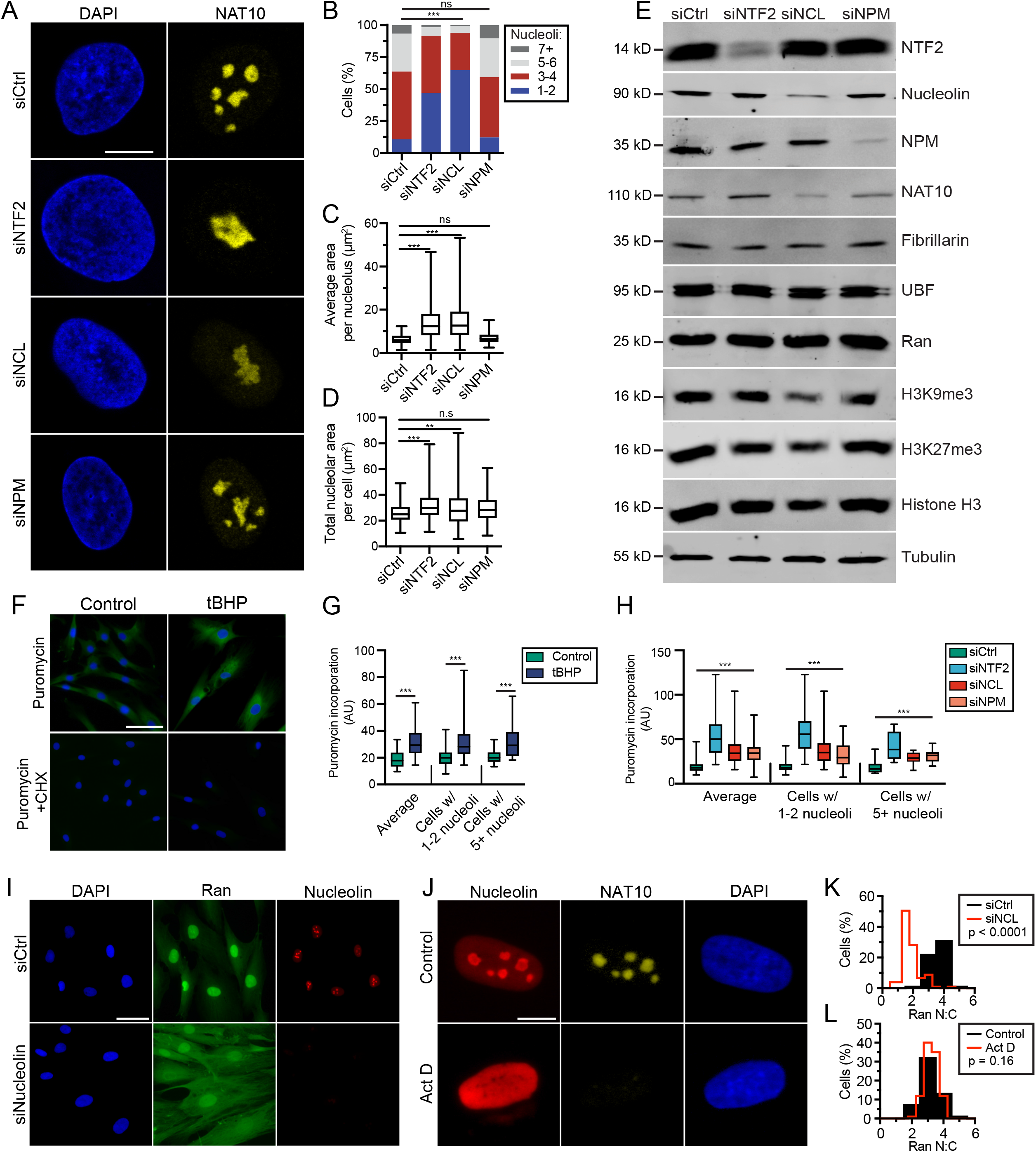
Nucleolin and nucleophosmin are critical determinants of nucleolar morphology and output. **(A)** Double label IF microscopy showing nucleolar morphology by NAT10 staining in normal human fibroblasts depleted of NTF2 (siNTF2), nucleolin (siNCL), and NPM (siNPM). Scale bar = 10 μm. **(B)** Number of nucleoli per cell quantified using NAT10 staining (n = 116). **(C)** Box and whiskers plot of average nucleolar size (n = 116). **(D)** Box and whiskers plot of total nucleolar size (n = 116). **(E)** Western blotting for nucleolar proteins, heterochromatin marks and tubulin (loading control) under indicated conditions. **(F-H)** SUnSET assay of fibroblast cells treated with tBHP, and depleted of NTF2, nucleolin, and NPM. **(F)** IF microscopy of cells treated with puromycin +/− cycloheximide (CHX). Scale bar = 100 μm. **(G)** Quantification of microscopy showing puromycin incorporation in cells treated with tBHP. (n = 110). **(H)** Quantification of microscopy showing puromycin incorporation in cells with depleted NTF2, nucleolin, and NPM (n=105). **(I)** Double label IF microscopy Ran and nucleolin in cells with depleted nucleolin. Scale bar = 60 μm **(J)**. Double label IF microscopy of Nucleolin and NAT10 in fibroblast cells treated with Actinomycin D (Act D). Scale bar = 10 μm. **(K)** Quantification of IF microscopy showing Ran N:C in cells with depleted nucleolin (n = 102). **(L)** Quantification of IF microscopy showing Ran N:C in cells treated with Actinomycin D (n = 100). In plots, boxes denote 25^th^ and 75^th^ percentile with median in the middle, and whiskers show minimal and maximal values. (ns, not significant, p > 0.05; *, p < 0.05; **, p < 0.001; ***, p < 0.0001, ANOVA).

In the course of characterizing the effects of nucleolin depletion on nucleolar morphology, we made the unexpected finding that nucleolin depletion is sufficient to disrupt the Ran gradient (**Fig. 4I, K**). Since NTF2 depletion alters nucleolar structure (**Fig. 4A-C**), our data could be interpreted as evidence that the Ran gradient operates both upstream and downstream of the nucleolus. The effect of nucleolin depletion on Ran distribution does not seem to be explained simply by a stress response triggered by loss of nucleolar structure, since disassembly of nucleoli with Actinomycin D had no obvious effect on Ran (**Fig. 4J, L**). In cells treated with Actinomycin D, nucleolin was localized to the nucleus, and cells displayed a steep Ran gradient (**Fig. S2**). It remains possible that the effect of nucleolin depletion on Ran reflects a nucleolus-independent effect, perhaps related to a nucleolin function in chromatin structure (Angelov et al., 2006).

It is not surprising that nucleolar structure is sensitive to the nuclear import of nucleolin and NPM, given these proteins are major components of the nucleolus and undergo continuous shuttling between the nucleus and the cytoplasm (Borer et al., 1989). For proteins undergoing rapid nucleocytoplasmic transport, even small changes in nuclear import efficiency could alter the steady state distribution. With these considerations in mind, we used an alternative strategy to examine the contribution of nuclear import to nucleolar structure that involved the Importin-β inhibitor, importazole (IPZ) (Soderholm et al., 2011). Treating normal human fibroblasts with IPZ (10 μM) for 24 hours increased nucleolar size and decreased nucleolar number to an extent that was comparable to that observed in cells with a disrupted Ran gradient, and the effect of IPZ on nucleoli was reversible upon drug washout (**Fig. 5A, B, C**). By IF microscopy, IPZ treatment reduced nucleolin and NPM levels in the nucleus, but within a few hours of drug removal, both proteins returned to the nucleus and concentrated in nucleoli (**Fig. 5D, E, F**). Restoration of nucleolar number occurred on a similar timescale as the import recovery **(Fig. 5G)**. Nucleolin and NPM protein did not change detectably during the IPZ treatment and washout **(Fig. 5H)**. Redistribution of nucleolin and NPM from the nucleus to the cytoplasm in IPZ treated cells was not obvious unless the images were captured with 5-fold longer exposure times (**Fig. S3**), presumably because of the ~10-fold dilution in the cytoplasm.

**Fig. 5.**
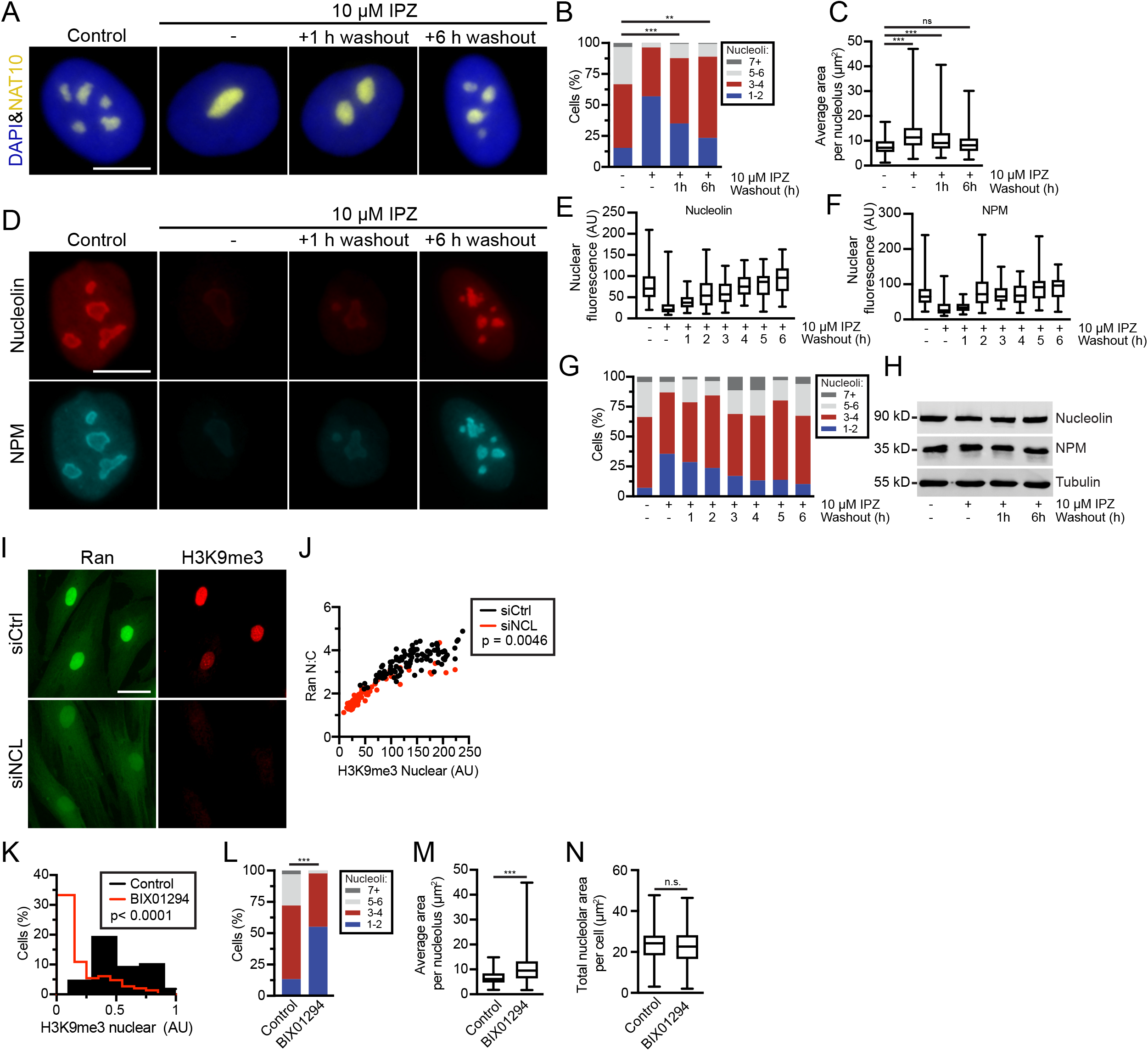
Nuclear transport and heterochromatin help specify nucleolar protein composition. **(A)** Double label IF microscopy showing NAT10 (yellow) merged with DAPI (blue) staining in normal human fibroblasts treated with importazole (IPZ) for 24 h, and allowed to recover for 1 and 6 h. Scale bar = 10 μm. **(B)** Number of nucleoli per cell quantified using NAT10 staining (n = 122). **(C)** Box and whiskers plot of average nucleolar size (n = 122). **(D)** Double label IF microscopy showing nucleolin and NPM in cells treated with IPZ for 24 h allowed to recover for up to 6 h. Scale bar = 10 μm. **(E, F)** Box and whiskers plot of total nuclear fluorescence of nucleolin and NPM under indicated conditions (n = 110). **(G)** Number of nucleoli per cell quantified using NAT10 staining (n = 110). **(H)** Western blotting for nucleolin, NPM, and tubulin (loading control) under indicated conditions. **(I)** Double label IF microscopy of ran and H3K9me3 in cells with depleted nucleolin. Scale bar = 40 μm. (**J**) Plot of Ran N:C versus H3K9me3 nuclear fluorescence from individual cells (n = 100). **(K)** Quantification of IF microscopy showing H3K9me3 nuclear fluorescence in cells treated with BIX01294 (n = 109). **(L)** Number of nucleoli per cell quantified using NAT10 staining (n = 104). **(M)** Box and whiskers plot of average nucleolar size (n = 104). **(N)** Box and whiskers plot of total nucleolar size (n = 104). In plots, boxes denote 25^th^ and 75^th^ percentile with median in the middle, and whiskers show minimal and maximal values. (ns, not significant, p > 0.05; *, p < 0.05; **, p < 0.001; ***, p < 0.0001, *t*-test, ANOVA)

Since nucleolin depletion disrupted the Ran gradient (**Fig. 4I, K**), and the Ran gradient is positively correlated with heterochromatin, we queried whether nucleolin influences the level of heterochromatin. Indeed, nucleolin depletion reduced heterchromatin levels, and and caused disruption of the Ran gradient (**Fig. 5I, J**). This observation together with the Ran-dependence of nucleolin import indicates an interdependence between the Ran gradient and nuclear nucleolin. We then tested for the interdependence between nucleolar structure and heterochromatin using the histone methyltransferase inhibitor BIX01294, which primarily targets G9a, and effectively reduces Histone H3K9me3 and Histone H3K27me3 levels (Kubicek et al., 2007). BIX01294 treatment was sufficient to increase nucleolar size and reduce nucleoli number, without affecting total nucleolar area (**Fig. 5K, L, M, N**). With BIX01294 treatment, there is a reduction in heterochromatin and a disruption of the Ran gradient (**Fig. S4A, B. C**). Taken together, the data suggests that heterochromatin levels, the Ran gradient, and ongoing nuclear transport is critical for maintaining nucleolar structure, and that perturbation of the system induces changes that resemble cellular phenotypes that occur in aging (**Fig. 6**).

**Fig. 6.**
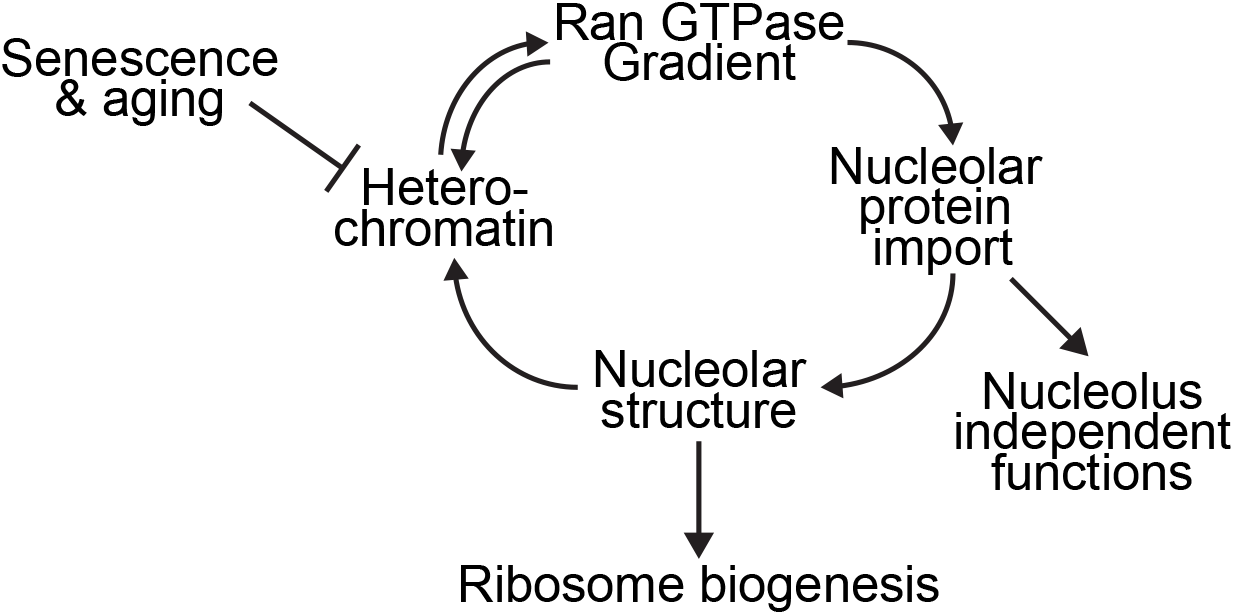
Model integrating the Ran GTPase with nucleolar structure. A Ran N:C protein gradient is required for efficient nuclear import of nucleolin, NPM, and other high molecular mass proteins with nuclear functions. A reduction in the Ran N:C gradient, which occurs in senescence and in premature aging, reduces import of nucleolin and NPM, and promotes an aging-related nucleolar phenotype. Nucleolin is also required to maintain heterochromatin levels, but it is not known if this is dependent or independent on the nucleolus. Disruption of the Ran gradient by depletion of its import factor, NTF2,is sufficient to reduce import of import of nucleolin and NPM and generate an aging-associated nucleolar phenotype.

## DISCUSSION

The steady state distribution of the Ran GTPase reflects a series of protein-protein interactions that control its GTPase cycle, nuclear import, and nuclear export. Perturbation of the Ran GTPase cycle via a temperature-sensitive mutant of its nucleotide exchange factor, RCC1, provided the first evidence that in mammalian cells, the nuclear distribution of Ran is linked to its GTPase cycle (Ohtsubo, Okazaki, & Nishimoto, 1989; Ren, Drivas, D’Eustachio, & Rush, 1993). It was subsequently determined that nuclear import of Ran from the cytoplasm is mediated by a single factor, NTF2 (Ribbeck et al., 1998; Smith et al., 1998; Steggerda et al., 2000). The fact that the Ran import factor, NTF2, is essential in *S. cerevisiae* emphasizes that maintaining the nuclear concentration of Ran is critical at the cellular level (Corbett & Silver, 1996; Paschal, Fritze, Guan, & Gerace, 1997).

The striking disruption of the Ran gradient in response to expression of the Progerin form of lamin A, and the resulting defect in nuclear import of large cargoes, led us to propose that the activity of the nuclear transport machinery is reduced in premature aging (Kelley et al., 2011; Snow et al., 2013). Here, we have extended these findings to show that Ran distribution and nuclear transport is compromised in cells that have undergone senescence. We found that the Ran gradient is disrupted in cells chemically induced to undergo senescence, and in cells that have undergone replicative senescence. A plausible, albeit simple explanation for these data, is that the Ran gradient responds to the level in heterochromatin (**Fig. 6**), one of the hallmarks of senescence and aging (Lopez-Otin et al., 2013). A mechanism that accounts for how Ran senses heterochromatin levels has not been elaborated, but could involve RCC1 since the nucleotide exchange reaction is stimulated by chromatin, and RCC1 binding is biased for heterochromatin (Dworak et al., 2019; Nemergut, Mizzen, Stukenberg, Allis, & Macara, 2001).

We found that Ran gradient disruption resulted in nuclear import defects for the nucleolar proteins, nucleolin and NPM, which are crucial components of the nucleolus (**Fig. 6**). Nucleolin and NPM localize to the dense fibrillar component and the granular component of the nucleolus, respectively, and participate in multiple steps of ribosome biogenesis. Nucleolin has more than 100 protein binding partners and is linked to a variety of RNA-based processes (Salvetti et al., 2016). NPM also has a complex network of protein interactions and participates in multiple nuclear pathways that include transcription (Colombo et al., 2011). Thus, nucleolin and NPM have important nucleolar-independent functions (Abdelmohsen & Gorospe, 2012; Mitrea et al., 2016). Our finding that nuclear localization of nucleolin and NPM proteins is highly sensitive to the Ran gradient is due to the large native sizes of these proteins, both of which rapidly shuttle between the nucleus and cytoplasm (Borer et al., 1989). In the case of nucleolin, we provided data in support of this hypothesis by using nucleolin NLS fusions with different-sized reporter proteins.

We also observed that Ran gradient disruption is sufficient to reduce the nucleoli number, and, slightly increase nucleolar size. This is interesting in light of the fact that nucleolar size and number are linked to aging in a variety of organisms (Tiku & Antebi, 2018). In C. elegans, small nucleoli are highly predictive for longevity, a finding that extends to flies, mice, and humans (Tiku et al., 2017). Since the Ran gradient is disrupted in senescent cells, we speculate that reduced nuclear import of key nucleolar proteins such as nucleolin and NPM modifies the self-organization of nucleoli (Misteli, 2001) in a manner that increases the size but limits the number; the outcome is enhanced ribosome output. Nucleolar phenotypes observed in senescent cells and in model organisms also occurs in fibroblasts from HGPS patients (Buchwalter & Hetzer, 2017). This observation places the structure of the nuclear lamina upstream of nucleoli (**Fig. 6**), and helps explains why in our experiments, LPV inhibition of lamin A processing resulted in the nucleolar phenotype detected in cells induced to undergo senescence. The nucleolar enlargement in HGPS cells, termed nucleolar expansion by the Hetzer group, was shown to be associated with increased translational capacity; this was also detected in normal fibroblasts from individuals of advanced age (Buchwalter & Hetzer, 2017). These observations, together with the finding that lamin B1 loss is biomarker for senescence (Freund, Laberge, Demaria, & Campisi, 2012), provide strong support for the view that the nuclear lamina undergoes protein composition changes during senescence that alter its structure, and these changes affect the structure and output of the nucleolus. Whether this reflects a direct effect on nuclear architecture, or an indirect effect through gene expression, remains to be determined. Alterations in nucleolar structure that enhance protein translation could be an adaptation that helps compensate for age-associated protein damage.

The nucleolus has long been recognized as a dynamic organelle, and this concept has been underscored by data showing that nucleolar proteins undergo phase separation on a rapid timescale, and that the process contributes to structure and possibly temporal regulation of ribosome biogenesis (Feric et al., 2016). Interestingly, one of the nucleolar proteins shown to undergo phase separation is NPM. Given the concentration dependence for phase separation of proteins, nuclear import is predicted to represent a limiting step for phase separation for nucleolar proteins that undergo continuous cycles of import-export. Consistent with this notion, inhibiting nuclear import with IPZ reduced nucleolar number and caused nucleolar enlargement. These nucleolar changes were associated with defective import of NPM and nucleolin. The effects of IPZ on nucleoli were reversible, indicating that on the time frame of our experiments, nucleoli with an enlarged, aging-type morphology retain the capacity for reversion to a normal morphology. Finally, given the evidence for nucleolar-independent functions of nucleolin and NPM, reduced Ran function and nuclear import may impact other nuclear events linked to senescence and aging.

## METHODS AND MATERIALS

### Cell culture

Normal human fibroblast cell line (AG08469, Coriell Institute for Medical Research) was maintained in Minimum Essential Medium (Gibco) and supplemented with 15% FBS (HyClone), 100 U/ml penicillin/streptomycin (Gibco), 1% MEM Vitamin Solution (Gibco), and 2 mM L-glutamine (Gibco). Cells used for experiments were between passage no. 10 and 16. HeLa cell line was grown in DMEM (Gibco) supplemented with 10% FBS (Atlanta Biologics) and 100 U/ml penicillin/streptomycin (Gibco). HEK293T were cultured in DMEM (Gibco) supplemented with 5% FBS (Atlanta Biologics), 1% non-essential amino acids (Gibco), 1% sodium pyruvate (Gibco) and 100 U/ml penicillin/streptomycin (Gibco). Cells were cultured at 37°C and 5% CO_2_.

### Induction of senescence

Senescence in normal human fibroblasts was induced using five treatments with 30 μM tert-butyl hydroperoxide (tBHP, Sigma) for 5 consecutive days (1 h/day) as previously published (Debacq-Chainiaux et al., 2009; Dumont et al., 2000). Cells were allowed to recover for two days before analysis. For replicative senescence induction, cells were passaged until exhaustion.

### Senescence-associated β-galactosidase (SA-β-gal) assay

Cells were incubated with fixation solution (2% formaldehyde and 0.2% glutaraldehyde in PBS) for 5 min. Then, cells were incubated with staining solution (40 mM citric acid/sodium phosphate buffer pH 6, 5 mM potassium hexacyanoferrate (II) trihydrate, 5 mM potassium hexacyanoferrate (III), 150 mM sodium chloride, 2 mM magnesium chloride and 1 mg/ml X-gal) at 37°C in CO_2_-free incubator for up to 16 hours overnight. Images were captured on Leica MZ16 with QImaging Micropublisher 5.0 RTV camera and QCapture v.2.8.1 software. For each condition, at least 100 cells were captured and quantified manually.

### Drug treatments

Lopinavir (LPV, Cayman Chemicals), was diluted in DMSO to 50 mM and used at 40 μM. Cells were treated with LPV or DMSO for 96 h. Actinomycin D (Act. D, Sigma Aldrich) was diluted in DMSO to 5 mM and used at 40 nM. Cells were treated with Actinomycin D or DMSO for 20 h. Importazole (IPZ, Selleckchem) was diluted in DMSO to 10 mM and used at 10 μM. Cells were treated with IPZ or DMSO for 24 h. After 24 h, cells were washed twice with PBS and incubated with fresh drug-free medium for further 1 and 6 h. BIX01294 (Cayman Chemicals) was diluted in DMSO to 10 mM and used at 2 μM. Cells were treated with BIX01294 or DMSO for 24 h.

### Cloning and constructs

The sequence of the bipartite nuclear localization signal (NLS) of nucleolin was based on published work(Creancier, Prats, Zanibellato, Amalric, & Bugler, 1993) and cNLS mapper. Synthetic oligos were designed to clone into pKGG (small cargo) and pRC/CMV (Invitrogen) with myc-pyruvate kinase (large cargo). NLS sequence was cloned into pKGG using BglII and EcoRI enzymes and the following oligos: forward 5’-GATCTAAACGAAAGAAGGAAATGGCCAAACAGAAAGCAGCTCCTGAAGCCAAGAA ACAGAAAGTGTAAG-3’, reverse 5’-AATTCTTACACTTTCTGTTTCTTGGCTTCAGGAGCTGCTTTCTGTTTGGCCATTTCCTTCTTTCGTTTA-3’. Cloning into pRC/CMV was done using NheI and ApaI using the following oligos: forward 5’-CTAGCGAAACGAAAGAAGGAAATGGCCAAACAGAAAGCAGCTCCTGAAGCCAAGAAACAGAAAGTGTAAGGGCC-3’, reverse 5’-CTTACACTTTCTGTTTCTTGGCTTCAGGAGCTGCTTTCTGTTTGGCCATTTCCTTCTTTCGTTTCG-3’.

### Plasmid and siRNA transfection

HeLa cells were grown in 6-well plates until reaching ~75% confluency. Cells were transfected using 1 μg DNA per well and ViaFect™ Transfection Reagent (Promega) in OptiMEM medium (1:3 DNA to reagent ratio). Cells were analyzed 24 h post-transfection. RNAi experiments were performed on fibroblast cell line using Lipofectamine™ RNAiMAX Transfection Reagent (Invitrogen), OptiMEM (Gibco) and the following siRNA at 10 nM: siNTF2 (Santa Cruz, sc-36105), siNucleolin (siNCL, Santa Cruz, sc-29230), and siNucleophosmin (siNPM, Santa Cruz, sc-29771), and siControl (Ambion, AM4635). Cells at 85-90% confluency were incubated with transfection mixture for 24 h. Then, cells were split and grown for another 96 h before analysis.

### Immunofluorescence Microscopy

For immunofluorescence experiments, cells were grown on glass coverslips. Cells were fixed with 3.75% formaldehyde in PBS for 10 min, and permeabilized with 0.2% Triton X-100 for 5 min. Then, cells were incubated with blocking solution (2% BSA, 2% FBS) for 30 min at RT or overnight at 4°C. Primary and secondary antibody solutions were prepared in blocking solution. Cells were incubated with primary antibodies for 2 h at RT and secondary antibodies for 1 h at RT. At the end, cells were briefly incubated with DAPI (1 μg/ml) and mounted onto a glass slide with Vectashield (Vector Laboratories). Images were acquired by laser scanning confocal microscopy Zeiss 710 LSM (Carl Zeiss) at 63×, 1.25 NA oil immersion objective with Zen software (Carl Zeiss), and Nikon Ni-U Eclipse Ni-U with Nikon DS-Ri1 camera, 10×, 20×, and 40× lenses with NIS Elements 4.13 software (Nikon).

### Immunoblotting

Immunoblotting samples were prepared by direct cell lysis with buffer (50 mM Tris-HCl pH 6.8, 2% SDS, 8% glycerol, 5% β-mercaptoethanol, and bromophenol blue). Samples were heated at 95°C for 5 min and sonicated.

### Antibodies

Primary antibodies used in this study: anti-Ran mouse mAb (BD Biosciences, #610341), anti-Ran rabbit pAb (Bethyl Laboratories, A304-297A), anti-Histone H3K9me3 rabbit pAb (Abcam, #8898), anti-Histone H3K27me3 rabbit pAb (Millipore, #07-449), anti-Histone H3 rabbit pAb (Abcam, ab1791), anti-NAT10 (B-4) mouse mAb (Santa Cruz, sc-271770), anti-Fibrillarin (C13C3) rabbit mAb (Cell Signaling, #2639), anti-Nucleolin (D4C7O) rabbit mAb (Cell Signaling, #14574), anti-Nucleophosmin (FC82291) mouse mAb (Santa Cruz, sc-56622), anti-UBF rabbit pAb (Sigma, HPA006385), anti-myc (9E10) mouse mAb (Abcam, ab32), anti-tubulin (1A2) mouse mAb (Sigma, T9028), anti-NTF2 (5A3) mouse mAb (made in laboratory, M. Steggerda), anti-puromycin (12D10) mouse mAb (Millipore Sigma, MABE343). Secondary antibodies: AffiniPure Donkey Anti-Mouse IgG conjugated with Alexa Fluor® 488 (Jackson Immunoresearch), AffiniPure Donkey Anti-Rabbit IgG (H+L) conjugated with Cyanine Cy™3 (Jackson Immunoresearch), Alexa Fluor 680® Donkey anti-Rabbit IgG (Invitrogen), Mouse IgG (H&L) Antibody DyLight™ 800 (Rockland).

### Quantification of Ran gradient and heterochromatin levels

Ran gradients were measured as a ratio of nuclear to cytoplasmic signal(Kelley & Paschal, 2019). Intensities were manually measured using ImageJ 1.52p (NIH). For each measurement, background signal was subtracted. Level of H3K9me3 in cells was measured by sampling the nuclear area of the cell based on DAPI staining. For each condition, at least 100 cells were counted.

### Quantification of nucleolar number and area

For quantification of nucleolar characteristics, an automatic pipeline was developed using Cell Profiler 3.15. Image analysis was based on DAPI staining and NAT10 immunofluorescence. Raw results were further cleaned up with R script to show the number of nucleoli per cell, area of every nucleolus in the nucleus, average nucleolar area per nucleus, and total nucleolar area per nucleus. For each condition, at least 100 nuclei were counted. The pipeline and script is available upon request.

### Gel filtration chromatography

HEK 293T were grown to confluency, harvested in ice-cold PBS with 1 mM PMSF and pelleted. Pellets were lysed for 20 min on ice in a PBS buffer with 100 mM NaCl, 0.5%Triton X-100, 2.5 mM EDTA, 2 mM DTT, and protease inhibitors, and then tip sonicated. Supernatant was clarified at 56000 rpm for 20 min at 4°C. Gel filtration chromatography was performed using an AKTA FPLC System with a Superose 6 Increase 10/300 GL column (GE Healthcare) in 50 mM NaPO4 pH 7.0, 150 mM NaCl with 1 mM DTT at a flow rate of 0.25 ml/min. The following standards were used (Sigma): Cytochrome C, Horse Heart (12.4 kD), Carbonic Anhydrase, Bovine Erythrocytes (29 kD) Albumin, Bovine Serum (66 kD), Alcohol Dehydrogenase, Yeast (150 kD) β-Amylase, Sweet Potato (200 kD), Apoferritin, Horse Spleen (443 kD), and Thyroglobulin, Bovine (669 kD).

### SUnSET assay

Cells on coverslips were treated with 1 μM puromycin (Gibco) alone or in combination with 100 μg/ml cycloheximide (CHX, Cayman Chemicals) for 15 min. Then, cells were processed as in immunofluorescence section. Incorporated puromycin in cells was detected using anti-puromycin antibody. To measure the puromycin signal, for each cell cytoplasmic signal was manually selected and quantified. Nucleoli in cells were detected by double labeling with nucleolin. In addition to puromycin signal, number of nucleoli was counted for every cell. For each condition, at least 100 cells were counted.

### Statistical analysis

The statistical analysis was conducted with GraphPad Prism 8 software. P values were determined using Student’s t test to measure the significance between two conditions, or one-way ANOVA with multiple comparisons when measuring significance between multiple conditions.

## ACKNOWLEDGEMENTS

We thank Chunsong Yang, Teddy Kamata, Roxanne Ruiz, Anna Mykytyn, and Jeff Smith for comments on the manuscript.

## AUTHORS’ CONTRIBUTIONS

B.P. and B.R. designed the study. B.R. performed the experiments and analyzed data. B.P. and B.R. wrote the manuscript.

## DATA AVAILABILITY STATEMENT

The data that support the findings of this study are available from the corresponding author upon reasonable request.

## SUPPLEMENTAL FIGURE LEGENDS

**Fig. S1.**
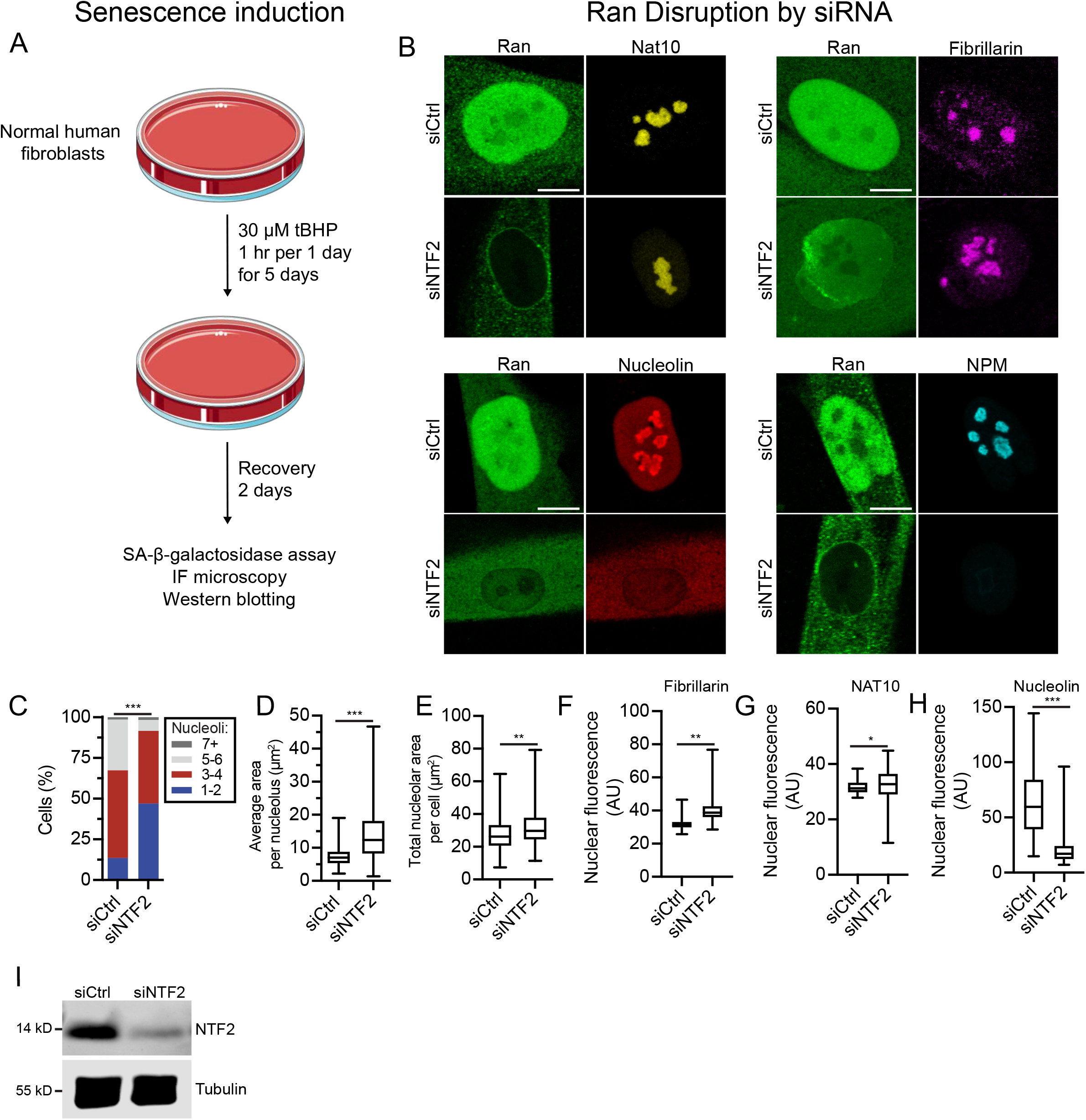
Senescence induction and Ran disruption in normal human fibroblasts. (A) Scheme for senescence induction, modeled after published methods (Debacq-Chainiaux et al., 2009; Dumont et al., 2000). Cell were treated with tBHP for 1 h on five consecutive days, allowed to recover for two days, and analyzed for senescence and protein localization. (B) IF microscopy for Ran and nucleolar proteins NAT10, nucleolin, fibrillarin, and NPM after depletion of NTF2. (C-H) Quantification of nucleolar area (NAT10 signal) and nuclear localization (fibrillarin, NAT10, nucleolin) in response to NTF2 depletion. (I) Immunoblot showing NTF2 protein levels after knockdown by siRNA using tubulin as a loading control.

**Fig. S2.**
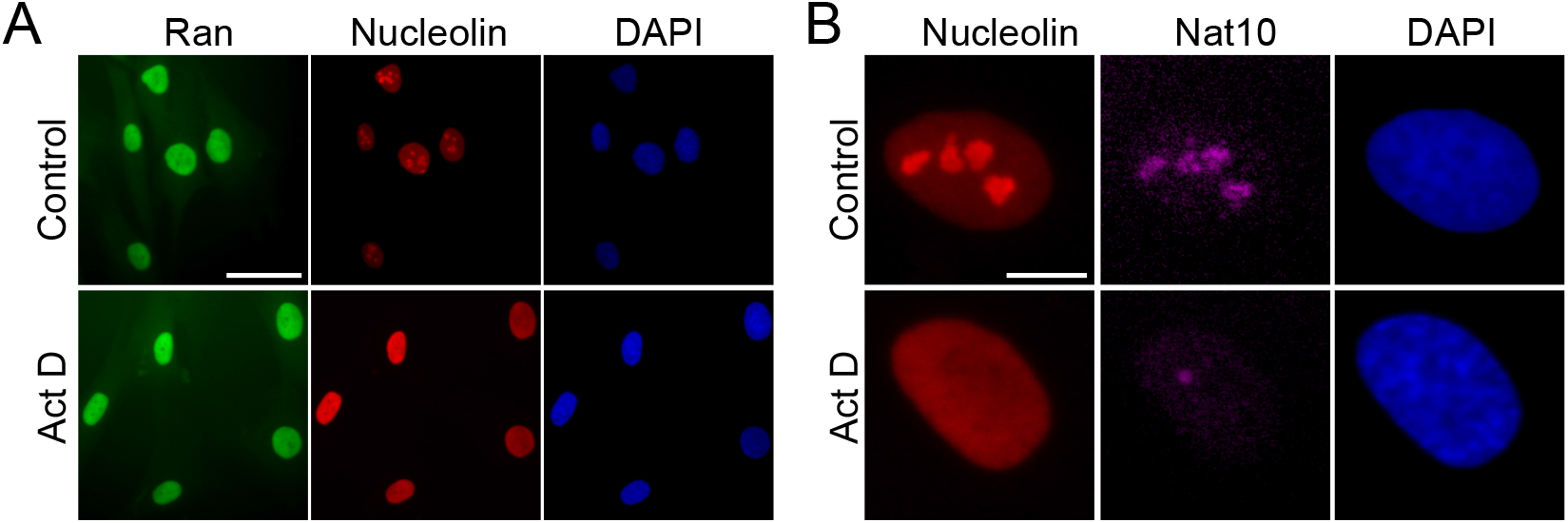
Actinomycin D treatment disrupts nucleolar structure and releases nucleolin into the nucleoplasm. (A) Double label IF microscopy for Ran and nucleolin in human fibroblasts treated with Actinomycin D (Act D). (B) Double label IF microscopy for nucleolin and NAT10 in human fibroblasts treated with Actinomycin D (Act D) shown at high magnification to more clearly show nucleoplasmic nucleolin.

**Fig. S3.**
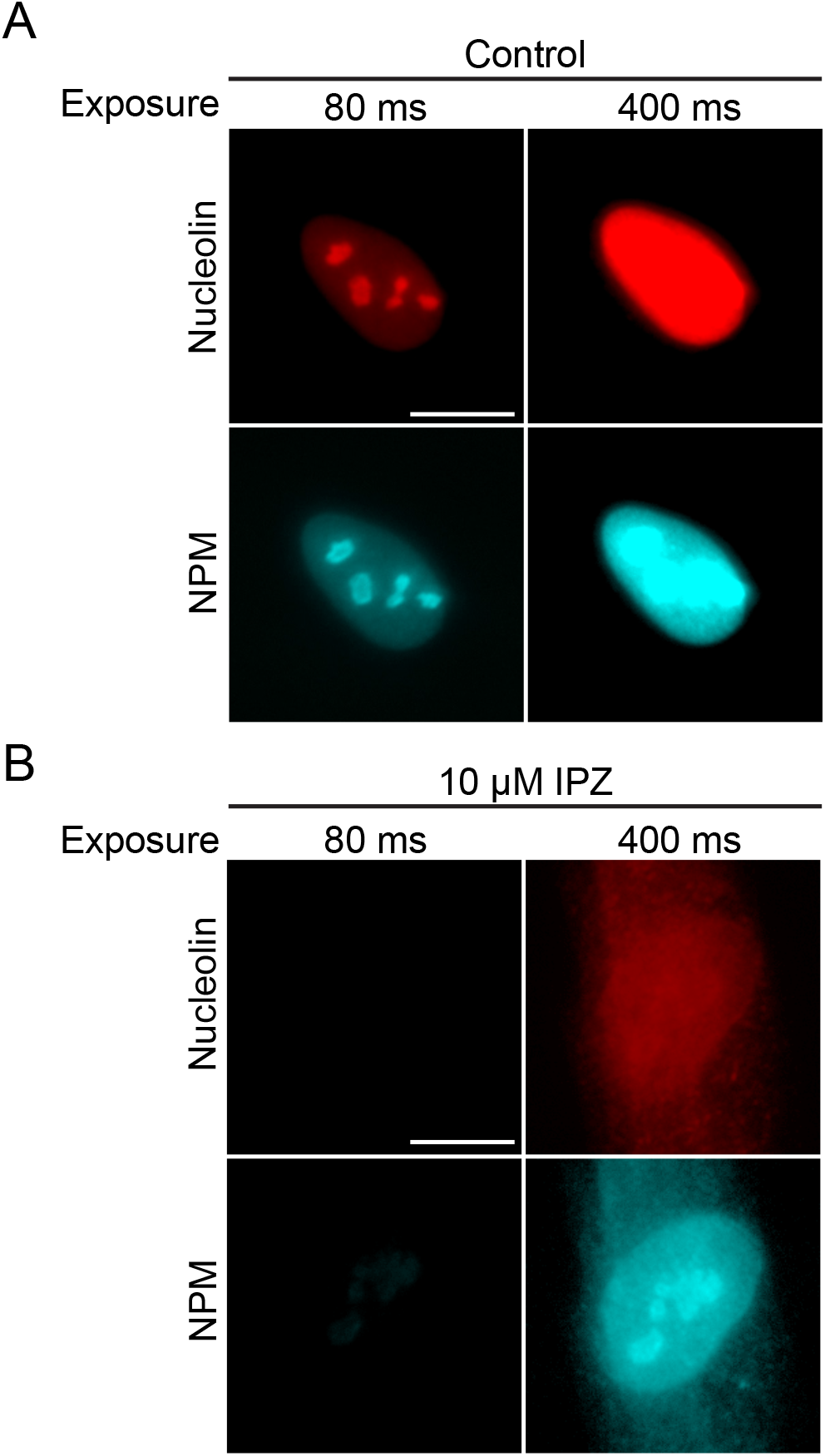
IPZ treatment reduces nuclear localization of nucleolin and nucleophosmin in human fibroblasts. (A) IF microscopy showing “normal” (80 millisecond) and “long” (400 millisecond) exposures of nucleolin and NPM in normal human. (B) Same as in (A) except the cells were treated with IPZ (10 μM for 24 h).

**Fig. S4.**
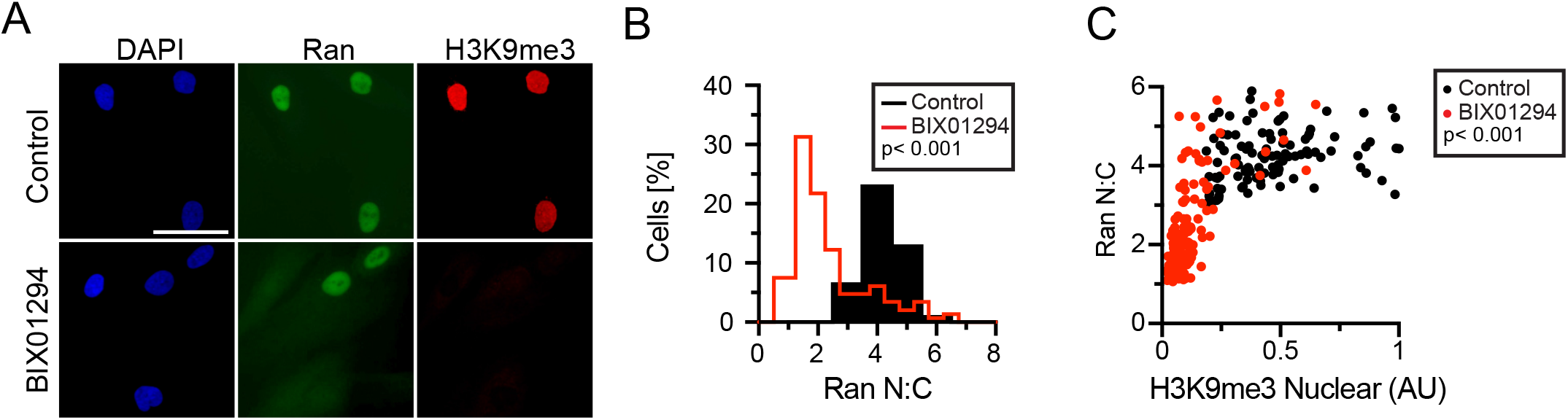
Reduction of heterochromatin levels in human fibroblasts using BIX01294. (A) Double label IF microscopy for Ran and Histone H3K9me3. (B, C) Quantification of Ran and Histone H3K9me3 levels in response to BIX01294.

